# Neutrophil microvesicles drive atherosclerosis by delivering *miR-155* to atheroprone endothelium

**DOI:** 10.1101/319392

**Authors:** Ingrid Gomez, Ben Ward, Celine Souilhol, Chiara Recarti, Mark Ariaans, Jessica Johnston, Amanda Burnett, Marwa Mahmoud, Le Anh Luong, Laura West, Merete Long, Sion Parry, Rachel Woods, Carl Hulston, Birke Benedikter, Rohit Bazaz, Sheila Francis, Endre Kiss-Toth, Marc van Zandvoort, Andreas Schober, Paul Hellewell, Paul C. Evans, Victoria Ridger

## Abstract

Neutrophils have been implicated in the pathogenesis of atherosclerosis, a lipid-driven disease of arteries, but they are seldom found in atherosclerotic plaques. To resolve this longstanding paradox, we investigated whether neutrophil-derived microvesicles may influence arterial pathophysiology. Clinical and pre-clinical studies revealed that levels of circulating neutrophil microvesicles were enhanced by exposure to a high fat diet, a known risk factor for atherosclerosis. Neutrophil microvesicles accumulated at disease-prone regions of arteries that are exposed to complex flow patterns, and they promoted vascular inflammation and atherosclerosis in a murine model. Using cultured endothelial cells exposed to disturbed flow, it was demonstrated that neutrophil microvesicles promoted inflammatory gene expression by delivering a microRNA (miR-155) that enhanced NF-κB activation. Similary, neutrophil microvesicles increased miR-155 and enhanced NF-κB at disease-prone sites of disturbed flow in arteries of mice. We conclude that delivery of microvesicles carrying miR-155 to disease-prone regions of arteries provides a novel mechanism by which neutrophils contribute to vascular inflammation and atherogenesis.

---

A causal role for neutrophils in atherosclerosis is now evident and these abundant leukocytes have been shown to play a part both in plaque development ^1,2^ and plaque erosion ^3^, as well as being implicated in plaque rupture ^4,5^. Increased levels of circulating neutrophils exacerbate atherosclerotic plaque formation in mice ^2^ and indirect evidence also links increased circulating leukocyte counts and infection with an increased risk of cardiovascular disease ^6,7^. Nevertheless, detection of neutrophils in lesions is rare, possibly due to their short life span, the rapid removal of senescent neutrophils by macrophages within the developing plaque, and/or the lack of a highly specific detection method (see ^8^ for review). In order to address this paradox, we have investigated whether neutrophils exacerbate vascular inflammation through the release of pro-inflammatory microvesicles (MVs), thus influencing atherosclerotic plaque formation without entering the vessel wall.

MVs are 0.1–1 µm vesicles released from the cell membrane in response to various stimuli or during apoptosis. Depending on the cell source, MVs vary in their size, content, and surface marker expression ^9^. While MVs are present in healthy individuals ^10^, increased levels of leukocyte MVs have been observed in patients with sepsis ^11^, acute vasculitis ^12^, and individuals with high risk of cardiovascular disease ^13^. Advances in our understanding of extracellular vesicle function have led to the discovery of a novel mechanism by which cells can communicate with each other through the transfer of vesicle cargo, such as noncoding RNA (e.g. microRNA (miRs)), to target cells ^14–17^. Neutrophil-derived microvesicles (NMVs) have been detected in human atherosclerotic plaques ^18^ but their role in plaque progression has not been studied.

Atherosclerotic plaque distribution is focal, with plaques developing at sites of disturbed flow, such as bifurcations, where adhesion molecules are highly expressed ^19,20^. Disturbed flow generates shear stress (mechanical drag) that is low and oscillatory at atheroprone sites, whereas shear is high at atheroprotected sites ^21,22^. Vascular inflammation is regulated by the transcription factor nuclear factor (NF)- κB ^23^, the expression of which is known to be enhanced in atheroprone regions ^24–26^ and induced by disturbed flow ^27^. Enhanced vascular inflammation leads to leukocyte recruitment to these sites, with the presence of monocytes within the vessel wall a characteristic of atherosclerotic plaque development.

Here we show that NMVs are released in response to high fat diet and preferentially adhere to sites prone to atherosclerotic plaque development. We also demonstrate NMVs contain miRs and are internalized by arterial endothelial cells. We investigated whether NMVs induce NF-κB expression, through delivery of cargo such as miRs, leading to enhanced endothelial inflammation, monocyte recruitment and atherosclerotic plaque development.

## Methods

### For detailed methods see supplemental online material

#### Ethics

For human studies, all experiments were approved by University of Sheffield Research Ethics Committee (reference SMBRER310) or by the Loughborough University Ethical Subcommittee (reference R13-P171). All subjects gave informed consent before the experimental procedures and possible risks were fully explained.

All procedures involving mice were approved by the University of Sheffield ethics committee and performed in accordance with the UK Home Office Animals (Scientific Procedures) Act 1986 under Project Licences 40/3562 and PF8E4D623. Where appropriate, age-matched animals were randomly assigned to treatment groups. Number rather than treatment group was used to label samples for subsequent blinded analysis.

#### High fat diet

##### Human experiments

Participants attended an initial pre-screening visit for assessment of baseline anthropometric characteristics and estimation of resting energy expenditure according to the calculations described by Mifflin *et al.* ^28^. From these procedures it was determined that a daily energy intake of 13474 ± 456 kJ was required to maintain energy balance. A week long, high fat diet intervention was carried out in order to increase daily energy intake by approximately 50% (19868 ± 759 kJ). Macronutrient intake was 333 ± 14 g [64%] fat, 188 ± 8 g [16%] protein, and 237 ± 8 g [20%] carbohydrate. Fasting venous blood samples were obtained in the morning before commencing the high fat diet and again after 7 days adherence. Blood samples were collected at least 12 hours after consuming the previous evening meal.

##### Mouse experiments

ApoE^-/-^ mice were fed chow or a high fat (21%) Western diet (8290; Special Diet Services, UK) for 6 - 20 weeks. Neutrophil depletion studies were carried out using i.p. anti-Ly6G antibody administration as previously described ^2,29^.

#### Neutrophil microvesicle isolation

NMV isolation was based on the method of Timár et al ^30^ with some modifications. In brief, human neutrophils were isolated from peripheral venous blood by density gradient separation. Mouse peripheral blood neutrophils were isolated by negative immunomagnetic separation as previously described ^31^. Isolated neutrophils were stimulated with the bacterial derived peptide *N*-formylmethionyl-leucyl-phenylalanine (fMLP 10 µmol/L; Sigma-Aldrich, MO) for 1 h (37 °C in 5% CO_2_). Neutrophils were then removed by spinning twice at 500 x *g* for 5 min and the supernatant collected. To remove residual fMLP, NMV suspensions were dialyzed using dialysis cassettes (ThermoFisher Scientific, MA). To pellet NMVs, the suspension was centrifuged at 20,000 x *g* for 30 min. NMV suspensions were tested for platelet contamination by flow cytometry using anti-human CD41 (BD Biosciences, UK) and for endotoxin contamination using Limulus Amebocyte lysate assay (Lonza, UK) and found to contain neither. NMV concentration from each isolation was quantified by flow cytometry. In order to detect NMVs, settings were standardized on an LSRII flow cytometer (BD Biosciences, UK) for forward (size) and side (granularity) scatter parameters using Megamix fluorescent calibration beads (BioCytex, France) according to the manufacturer’s instructions (Supplemental Figure 1A-C). NMVs were quantified using Sphero(tm)AccuCount beads (Saxon Europe, UK) as described in the Supplemental Material. For experiments where fluorescently-labelled NMVs were required, PKH26 or PKH67 fluorescent cell linker kits for general cell membrane labelling were used (Sigma, UK) according to the manufacturer’s instructions. We have provided transmission electron micrographs (Figure 1A wide field to show heterogeneity and B close up of single vesicle) as well as size data from Tunable Resistive Pulse Sensing (Figure 1C; mode size 280 ± 16.6 nm) to demonstrate the content of our microvesicle pellet prepared from isolated neutrophils. This is in accordance with the current recommendations of the International Society for Extracellular Vesicles ^32^ for characterization of single vesicles. In addition, we also investigated the expression of proteins that neutrophil-derived microvesicles may inherit from their origin cell, as suggested by the International Society of Extracellular Vesicles. Methods are described in detail in the Supplemental Material.

**Figure 1.**
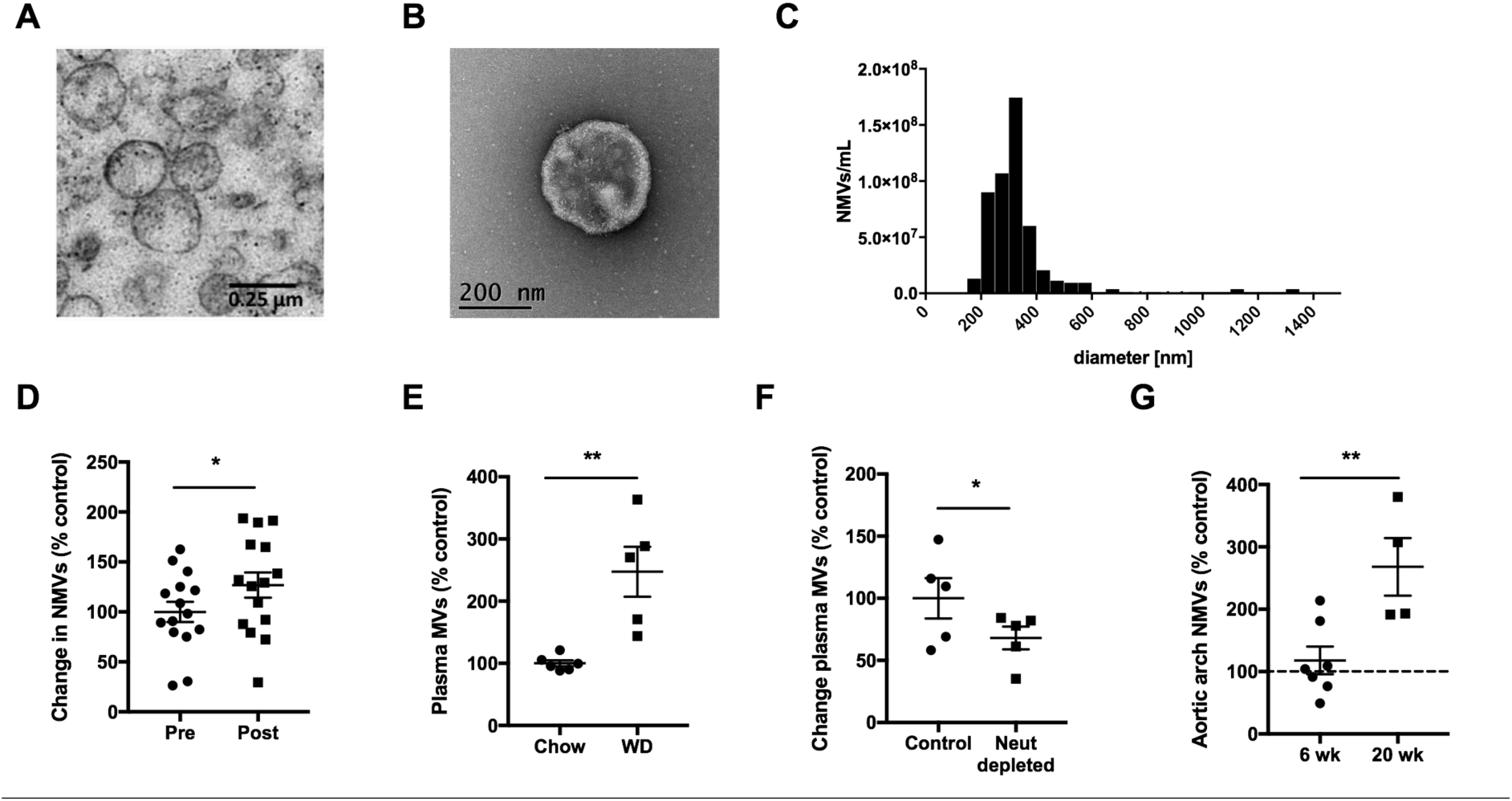
Hypercholesterolemia promotes the release of neutrophil microvesicles, which accumulate in the vessel wall. (**A-C**) NMVs were prepared from stimulated human neutrophils. (**A**) Transmission electron micrograph of an NMV pellet. (**B**) Transmission electron micrograph of a negatively-stained NMV sample on carbon-coated copper grids. Magnification of both micrographs 28,500x. (**C**) Representative histogram showing size distribution of NMVs analysed using Tunable Resistive Pulse Sensing. (**D**) NMVs were detected in human plasma samples before and after high fat diet using flow cytometry by staining with FITC-anti-CD66b (n = 15). (**E**) Total plasma MVs in *ApoE*^-/-^ mice fed chow or high fat diet (n = 5 - 6) and (**F**) in *ApoE*^-/-^ mice fed high fat diet with and without neutrophil depletion (n = 5) were quantified by flow cytometry. Numbers were normalised to the mean of control samples (filled circles, **D - F**). (**G**) NMVs were detected in aortic arch homogenates by staining with FITC-anti-mouse Ly6G. The accumulation of NMVs in the aortic arch of *ApoE*^-/-^ mice on Western diet for 6 – 20 weeks was compared to that of *ApoE*^-/-^ mice on chow (dotted line) using flow cytometry (n = 4 - 7). Data are presented as mean ± SEM and statistical significance evaluated using a paired (**D**) or unpaired (**E - G**) *t*-test.

#### Preparation of mouse aortic arch homogenates

Aortic arch homogenates were prepared from *ApoE*^-/-^ mice fed chow diet for 20 weeks, or Western diet for 6 weeks or 20 weeks based on the method described by ^18^.Mice were culled by i.p. injection of pentobarbital overdose and aortae perfused *in situ* with ice cold PBS. Aortic arches were dissected in ice cold PBS, rinsed in DMEM and minced thoroughly using fine scissors and forceps in 1ml of DMEM. After centrifugation at 400 x *g* for 15 min to remove contaminating cells, the supernatant was transferred and was further centrifuged at 5000 x *g* for 5 min to eliminate cellular debris. The homogenate was then analysed by flow cytometry.

#### Multicolor flow cytometry analysis of microvesicles

MV levels in platelet poor plasma or tissue homogenates were assessed by multicolor flow cytometry to detect specific surface markers derived from the parent cell (details are given in Supplemental Material). MVs were detected and quantified as described above. The number of events in the positive gate for each marker was quantified using FlowJo analysis software (Tree star Inc, Ashland, OR) and the total number of MVs in each subpopulation calculated from the total. The gating strategy for analysis of human plasma MVs is shown in Supplemental Figure 1E and for mouse aortic arch homogenate in Supplemental Figure 2.

#### Neutrophil microvesicle adhesion and internalization *in vivo*

To assess NMV adhesion and internalization, saline (150 µl) or fluorescently-labelled NMVs (4 × 10^6^ in 150µl) were injected via the tail vein into *ApoE*^-/-^ mice fed a Western diet for 6 weeks. After 2 h, mice were culled by i.p. injection of pentobarbital overdose and *en face* immunostaining of the mouse aortic arch was carried out as previously described ^25,27,33^ and described in detail in the Supplemental Material.

#### Human monocyte isolation

In experiments where NMVs and monocytes were both used, cells were isolated from the same donors. Following density gradient separation, peripheral blood mononuclear cells were harvested and monocytes isolated using negative immunomagnetic separation according to the manufacturer’s instructions (Monocyte Isolation Kit II, Miltenyi Biotec, Germany).

#### Human coronary artery endothelial cell (HCAEC)

Primary human coronary artery endothelial cells (HCAEC) were from Promocell (Germany). Endothelial cell preconditioning to controlled flow conditions was performed using the Ibidi pump system (Ibidi, Germany). HCAECs (1.7 × 10^5^) were seeded onto gelatin coated, 0.4 µm deep flow chambers (Ibidi µ-slide I^0.4^ Leur, Ibidi, Germany) and incubated for 2 h at 37 °C and 5% CO_2_ to allow adhesion. HCAECs were cultured under high unidirectional shear stress (HSS; 13 dyne/cm^2^) or low oscillatory shear stress (OSS; 4 dyne/cm^2^, oscillating at 1 Hz) for 72 h.

#### Neutrophil microvesicle adhesion *in vitro*

HCAEC were cultured under shear stress for 72 h as described above. Fresh complete growth medium containing fluorescently-labelled NMVs (1 × 10^3^/µL) was added to each Ibidi µ-slide for 2 h at 37 °C and 5% CO_2_. Following incubation, the medium was removed and cells gently washed three times with PBS to remove residual NMVs. Phase-contrast and fluorescent images were taken using the 20X lens of a wide-field microscope (Leica, DM14000B) and the mean number of fluorescent NMVs in 6 fields of view per sample was calculated.

To assess NMV adhesion to monocytes, fluorescently-labelled NMVs (1 × 10^3^/µL) were incubated with 2 × 10^5^ monocytes for 2 h at 37 °C. Unbound NMVs were removed by centrifugation (400 x *g* for 6 min) and the cells washed. NMV adhesion was analysed using an LSRII flow cytometer and data analysed for changes in mean fluorescence intensity using FACSDiva acquisition software.

#### Adhesion of monocytes to HCAEC under flow *in vitro*

HCAEC were cultured under OSS for 72 h as described above. Unlabelled NMVs (2 × 10^3^/µL) were perfused over the cells under OSS for 2 or 4 h. The media was removed and replaced with media containing fluorescently-labelled monocytes (1 × 10^3^/µL) and this was perfused over the HCAEC for 2 h under OSS. Slides were washed gently to remove non-adherent monocytes and fixed in paraformaldehyde (4% w/v). Phase-contrast and fluorescent images were taken using the 10X objective of a wide-field microscope (Leica, DM14000B) to detect adherent monocytes. An average of 15 images were analysed per slide and used to calculate the number of adherent monocytes per field of view per sample.

#### Monocyte transendothelial migration *in vitro*

HCAEC were cultured on transwell inserts as previously described ^34^. Monolayer integrity was checked using FITC-BSA, as previously described ^35^, and found to retain > 80% FITC-BSA in the upper chamber of the plate after 120 min. HCAEC were incubated ± NMVs (1 × 10^3^/µL) for 30 min. Subsequently, 2 × 10^5^ monocytes were added to the upper chamber and the number of monocytes that had migrated into the lower chamber in response to CCL2 (5 nmol/L) was counted after 90 min. To determine the role of HCAEC, the experiment was repeated in the absence of HCAEC. To determine the effects of CD18 inhibition, NMVs were treated with anti-CD18 (6 µg/10^6^ MVs; 6.5E gifted from M. Robinson, SLH Celltech Group, UK) or isotype control for 20 min at room temperature. Unbound antibody was removed by washing twice and the transendothelial migration experiment was repeated.

#### Flow cytometry analysis of surface molecule expression

MVs, NMVs, HCAEC and monocytes were stained with fluorescently-conjugated antibodies and an LSRII flow cytometer was used to detect the MV/cell population using forward (size) and side (granularity) scatter parameters. Changes in mean fluorescence intensity were analysed using FACSDiva acquisition software. See Supplemental Material for antibody details.

#### Cytometric bead array

HCAEC (3 × 10^4^) were cultured 72 h prior to the experiment in a 24 well plate. Cells were incubated ± NMVs (1 × 10^3^/µL) and media collected at 2 h and 4 h. CCL2, IL-6 and CXCL8 levels were assessed using a cytometric bead array (BD Biosciences, UK), a flow cytometry application that allows quantification of multiple proteins simultaneously.

#### Western blot analysis

HCAEC (3 × 10^4^) were cultured 72 h prior to the experiment in a 24 well plate. Cells were incubated ± NMVs (1 × 10^3^/µL) for 2 h. HCAEC were then washed, lysed, and centrifuged to remove any cellular debris. Western blotting was performed and membranes were imaged and the optical density analysed using a LI-COR C-DiGit^®^ Blot scanner (LI-COR Biosciences, NE). Detail provided in the Supplemental Material.

#### RNA extraction and real-time quantitative reverse-transcription polymerase chain reaction

Sample preparation is described in the Supplemental Material. RNA and miR were extracted using RNA isolation kit (Bioline, UK) and Pure Link microRNA isolation kit (Invitrogen, CA) respectively and reverse transcription polymerase chain reaction (RT-PCR) performed with 0.1 µg of RNA or miR using a OneStep kit (Qiagen, Germany); quantitative PCR (qPCR) was performed using SYBRgreen (Sigma-Aldrich, MO) for RNA analysis and Taqman (Eurogentec, Belgium) for miRNA. Relative gene expression was calculated comparing the number of cycles required to produce threshold quantities of product and calculated using the ΔΔCT method. β actin was used as housekeeping gene to normalize regulation of expression.

miR copy number was determined by spiking samples with known amounts of custom RNA oligonucelotides (Sigma Aldrich, MO) corresponding to the mature miR sequences. RNA oligos were serially-diluted in RNAse free water and amplified by qPCR. A standard curve was generated from the C_t_ value equivalent to the known amount of RNA oligo in each Taqman qPCR reaction as previously described ^36 37^ and the copy number was calculated using the molar mass of the synthetic RNA oligonucleotides (Supplemental Figure 3 A – B).

#### Microscopy analysis of neutrophil microvesicle internalization *in vitro*

Live cell imaging was performed in HCAEC. Cells were seeded on glass bottomed µ-slides (Ibidi) and cultured for 24h. SiR-Actin (Spirochrome, Switzerland) and Hoechst (ThermoFisher Scientific, MA) staining was performed to detect F-actin and nuclei in living cells respectively. Cells were then incubated with fluorescently-labelled NMVs and confocal live cell imaging performed on a Leica TCS SP8 imaging platform (Leica, Germany) equipped with an incubator, allowing live cell imaging at 37 °C and at 5% CO_2_. Samples were observed using a 100X oil immersion objective (HCX PL APO 100X/1,40 oil) and excited using sequential scanning, first at 405 nm and 652 nm for Hoechst and SiR-actin, respectively and at 490 nm for PKH67. HyD hybrid detectors were used to detect three spectral regions: 415-496 nm (Hoechst), 662-749 nm (SiR-actin), and 500-550 nm (PKH67). Z-stacks were performed with a z-step size of 0.20 µm. Images were analysed using ImageJ (1.49V, NIH) and Amira 6 software (ThermoFisher Scientific, MA).

#### Flow cytometry quantification of neutrophil microvesicle internalization *in vitro*

In order to quantify NMV internalization, HCAEC (5 × 10^4^) were seeded onto 24 well tissue culture plates and cultured overnight. The next day, fluorescently-labelled NMVs (0.4 × 10^3^/µL) were added and cells incubated for 2 h at 4°C, room temperature or 37°C (as specified in the corresponding figure legends). Cells were then washed to remove excess NMVs and detached using a trypsin/EDTA solution. Cells were washed and, immediately prior to flow cytometry analysis, Trypan blue (1 mg/mL) was added to each sample in order quench fluorescence of residual surface bound NMVs as previously described ^38,39^, thus ensuring any fluorescent signal detected was from internalized NMVs only. Data were analysed for mean fluorescence intensity using an LSRII flow cytometer using FACSDiva acquisition software.

#### Transfection with antagomir

HCAECs were transfected with antagomir designed to block *miR-155* regulation of target gene expression according to manufacturer’s instructions (Creative Biogene, NY). See Supplemental Material for details.

#### Atherosclerotic plaque analysis

6 week old *ApoE*^-/-^ mice were fed a Western diet for 6 weeks. During this period, 4 × 10^6^ NMVs, or an equivalent volume of saline, were injected twice weekly via the tail vein. *En face* staining with Oil-Red-O was then performed and lesion coverage in aortae was analysed as previously described ^40^. Monocyte/macrophage accumulation was assessed by *en face* staining with anti-MAC-3. See Supplemental Material for details.

#### Statistical analysis

Results are presented as mean ± SEM throughout. Statistical analyses were performed using GraphPad Prism version 7.00 (GraphPad Software, CA). Data was analysed using paired or unpaired *t*-tests, one-way ANOVA followed by Tukey’s post hoc test for multiple comparisons or Dunnett’s post hoc test to compare to control values, or two-way ANOVA followed by Tukey’s or Bonferonni’s post hoc test for multiple comparisons. *P* values of less than 0.05 were considered significant. For *in vitro* experiments, n numbers relate to different donors for both HCAEC and neutrophils/monocytes. In experiments where percentages are shown, data are expressed as the percentage of the mean of the control samples.

The datasets generated during and/or analysed during the current study are available from the corresponding author on reasonable request.

## Results

### Plasma neutrophil microvesicle levels are elevated in response to proatherogenic diet

We determined whether exposure to a high fat diet in healthy human subjects affected circulating levels of NMVs. The energy intake and diet composition is described in the online methods and an example of the typical daily food intake is shown in Supplemental Table 1. Flow cytometry analysis revealed that human plasma NMV levels were significantly increased after 1 week of high fat feeding (approximately 27% increase, Figure 1D) indicating that a high fat diet induced increased curculating NMV levels. Analysis of markers of different cellular origins revealed that MVs derived from neutrophils, platelets and monocytes were significantly increased after high fat feeding (Supplemental Tables 2 and 3 and Supplemental Figure 4). We also found elevated levels of total plasma MVs in *ApoE*^-/-^ mice on high fat diet compared to chow (Figure 1E) and neutrophil depletion for 48 h prior to plasma collection significantly reduced circulating MV levels compared to control (approximately 32%, Figure 1F). Taken together, these findings provide evidence that NMV are produced *in vivo* in response to a proatherogenic diet.

### Neutrophil microvesicles preferentially adhere to atheroprone regions in hypercholesterolemia

Having determined that high fat diet induced production of NMVs, we investigated whether these endogenously released NMVs were detectable in the vessel wall. Flow cytometry analysis of aortic arch homogenates from *ApoE*^-/-^ mice fed chow or a Western diet revealed that NMVs accumulated in the vessel wall at 20 weeks compared to 6 weeks (Figure 1G). Of note, significantly more platelet and monocyte derived MVs were also detected in the homogenates (Supplemental Table 4 and Supplemental Figure 5) at 20 weeks. In order to investigate the mechanisms by which NMVs accumulate in the vessel wall, we determined whether NMVs were able to adhere to arteries *in vivo*. Fluorescently-labelled NMVs (4 × 10^6^) or saline were injected via the tail vein into *ApoE*^-/-^ mice that had been fed a Western diet for 6 weeks. Using *en face* confocal microscopy, fluorescently-labelled NMVs were detected in atheroprone regions (inner curvature of aortic arch; Figure 2C and D) after 2 h but significantly fewer were detected at the atheroprotected regions (outer curvature of aortic arch; Figure 2A and B; quantified in Figure 2G). No fluorescence was detected in *ApoE*^-/-^ mice that were injected with saline (Figure 2E and F). Thus, we conclude that NMVs adhere *preferentially* to atheroprone sites within arteries in conditions of hypercholesterolemia.

**Figure 2.**
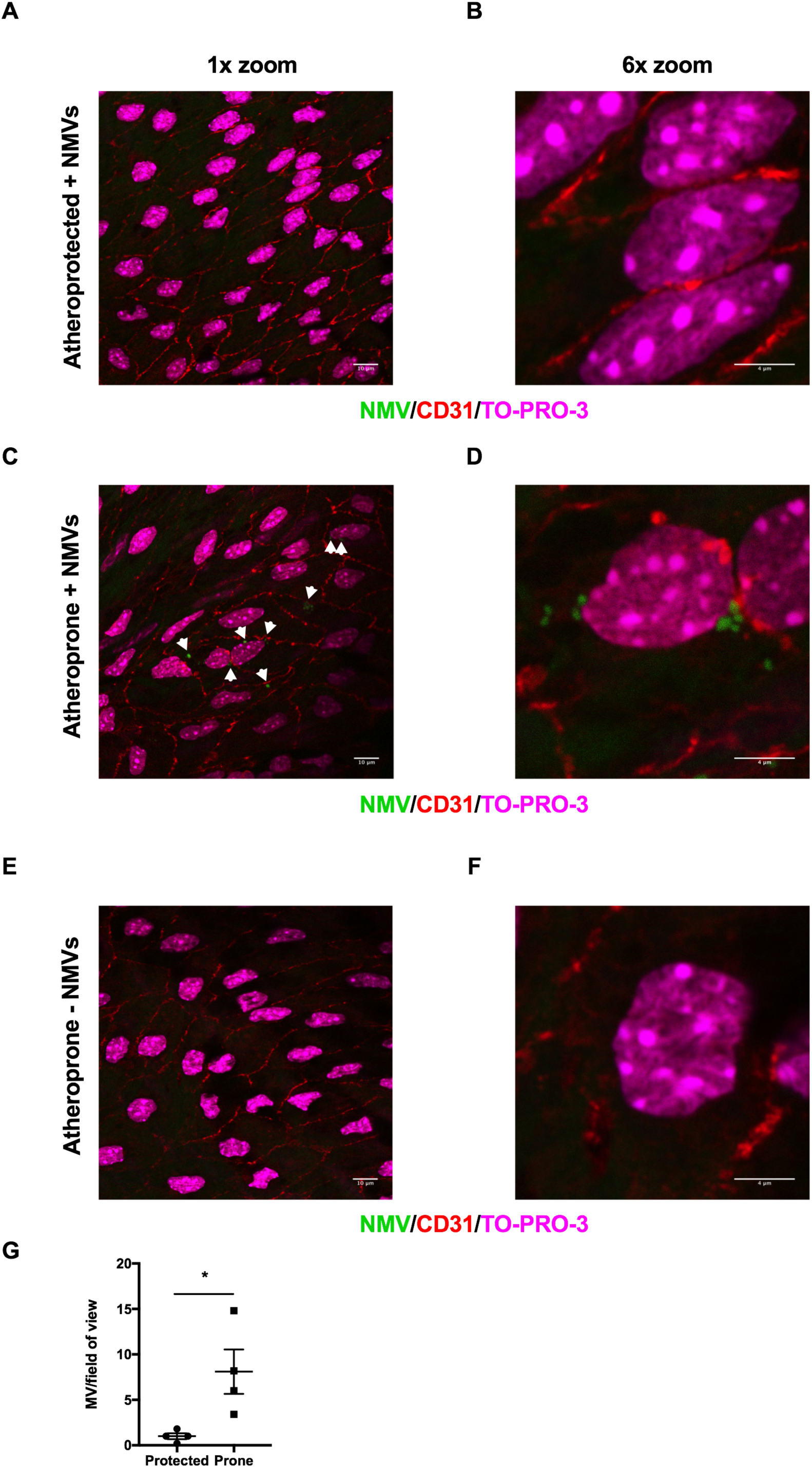
Neutrophil microvesicles preferentially adhere to atheroprone regions *in vivo*. Fluorescently-labelled NMVs (green; **A - D**) or saline (**E - F**) were injected via the tail vein into *ApoE*^-/-^ mice that had been fed a Western diet for 6 weeks. After 2 h, mice were culled and *en face* immunostaining of the mouse aortic was performed. Representative *en face* images of NMV adhesion in atheroprotected (outer curvature, **A - B**) and atheroprone (inner curvature **C - D**) regions of the aorta, visualized by confocal fluorescence microscopy. Representative *en face* images of atheroprone regions of saline injected mouse are also shown for comparison (**E - F**). Endothelial cells were identified by staining with anti-CD31 antibody (red) and cell nuclei were identified using TO-PRO Iodide (magenta). Outer and inner curvature of the ascending aorta were identified by anatomical landmarks and confirmed by characterising the phenotype of endothelial cells; those at the outer curvature were aligned, elongated and uniform – a characteristic of cells under high shear, whereas cells in the inner curvature had a disorganized appearance. Samples were visualised using a 100x objective at 1x zoom (**A, C, E**; scale bar = 10 µm) and at 6x zoom (**B, D, F**; scale bar = 4 µm). (**G**) Quantification of adherent NMVs presented as mean ± SEM (n = 4) and statistical significance evaluated using a paired t-test test. * *P* < 0.05.

### Oscillatory shear stress promotes neutrophil microvesicle adhesion to arterial endothelial cells

In order to investigate the mechanism of preferential adhesion of NMVs to atheroprone endothelium, we carried out *in vitro* adhesion experiments using HCAEC cultured under different flow parameters. We used oscillatory shear stress (OSS) to model flow at atheroprone regions and high shear stress (HSS) to model the flow found at atheroprotected areas. As with our *in vivo* data, we found that more NMVs adhered to HCAEC that had been cultured under OSS compared to HSS (Figure 3A – B). As NMVs have previously been shown to adhere to endothelial cells via a CD18-dependent mechanism ^41^, we confirmed the presence of adhesion molecules on the surface of NMVs as previously described ^35,41,42^ (Supplemental Figure 6A). We then hypothesized that the preferential adhesion to atheroprone regions may be due to alterations in endothelial expression of the CD18 counter-receptor, intercellular adhesion molecule-1 (ICAM-1). Indeed, HCAEC cultured under oscillatory flow expressed higher levels of ICAM-1 on their surface compared to cells exposed to high shear (Figure 3C). Consequently, pretreatment of HCAEC cultured under oscillatory shear stress with anti-ICAM-1 antibody significantly inhibited NMV adhesion (Figure 3D) indicating the preferential adhesion of NMVs to HCAEC exposed to OSS was via an ICAM-1-dependent mechanism. Although PSGL-1 was detected on the surface of NMVs (Supplemental Figure 6A), HCAEC levels of P-selectin were found to be unchanged by shear stress (Supplemental Figure 6B) and therefore not likely to mediate the preferential adhesion of NMVs to atheroprone regions.

**Figure 3.**
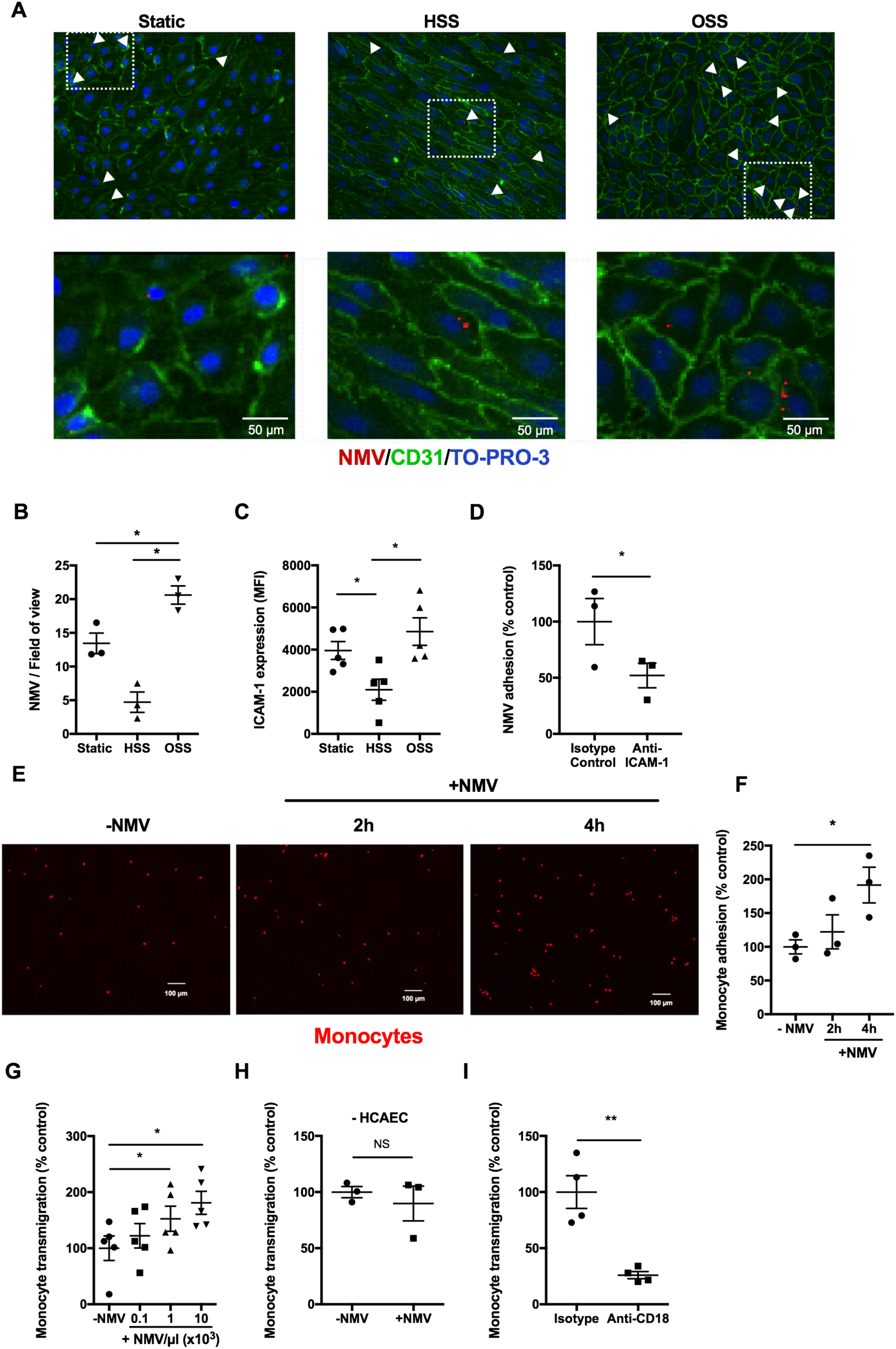
Adhesion of neutrophil microvesicles to HCAEC and subsequent effects on monocyte transendothelial migration. (**A - B**) Fluorescence microscopy of HCAEC cultured for 72 h under static conditions, high laminar (HSS) or low oscillating shear stress (OSS), incubated with fluorescently-labelled NMVs (red) for 2 h was performed. Cells were co-stained for CD31 (green) and TO-PRO-3 (blue). (**A**) Representative images of cells under each condition are shown. Lower panel shows higher magnification images of the areas outlined by the dotted rectangle in the upper panel. Scale bar = 50 µm. (**B**) NMV adhesion was quantified using fluorescence microscopy and the mean number of fluorescent NMVs in 6 fields of view per sample was calculated (n = 3). (**C**) HCAEC were cultured for 72 h under static, HSS or OSS followed by incubation with fluorescently labelled anti-ICAM-1. Changes in surface ICAM-1 expression were analysed by flow cytometry and are displayed as the mean fluorescence intensity (MFI) (n = 5). (**D**) HCAEC cultured for 72 h under OSS were incubated with anti-ICAM-1 or isotype control antibody (100 µg/mL) for 1 h prior to addition of fluorescently-labelled NMVs and adhesion measured using fluorescence microscopy (n = 3). Data are expressed as a percentage of the mean of the isotype control sample. (**E**) HCAEC were condition under OSS for 72 h. Non-fluorescent NMVs were perfused over conditioned HCAEC for 2 h. The media was then replaced with perfusion media containing fluorescently-labelled monocytes (red) for 2 - 4 h and (**F**) the number of adherent monocytes assessed by fluorescence microscopy (n = 3). (**G - I**) HCAEC were cultured on transwell inserts and incubated with (+NMV) or without (-NMV) NMVs for 30 min followed by addition of monocytes. The number of monocytes that transmigrated into the lower chamber in response to CCL2 (5 nmol/L) was measured after 90 min (n = 5). (**H**) Monocyte transmigration was repeated in the absence of the endothelial monolayer (n = 3). (**I**) NMVs were treated with anti-CD18 or isotype control for 20 min, washed and the transendothelial migration experiment repeated (n = 4). (**F - I**) Data are expressed as a percentage of the mean of control samples (-NMV/isotype) and are presented as mean ± SEM. Statistical significance was evaluated using a paired *t*-test (**D, H, I**) or one-way ANOVA followed by Tukey’s (**B, C**) or Dunnett’s (**F, G**) post hoc test. * *P* < 0.05, ** *P* < 0.01.

### Neutrophil microvesicles enhance monocyte transendothelial migration

To determine the functional consequences of NMV interaction with arterial endothelial cells, we investigated their potential effects on pathophysiological processes underpinning vascular inflammation and atherogenesis. *In vitro* experiments showed that the presence of NMVs significantly enhanced monocyte adhesion to HCAEC under OSS (Figure 3E - F). Furthermore, monocyte transendothelial migration to CCL2 was increased in the presence of NMVs in a manner that was dependent on the number of NMV present (Figure 3G). Crucially, NMVs released from unstimulated neutrophils were unable to induce this increase in monocyte transmigration (Supplemental Figure 7A), suggesting that this response is dependent on the properties of NMVs released from stimulated neutrophils. The effect was, however, dependent on the presence of endothelial cells, as NMVs did not influence monocyte migration in the absence of HCAEC (Figure 3H). Pre-blocking CD18 on the surface of NMVs significantly reduced monocyte transendothelial migration (Figure 3I). We noted that relatively few fluorescently-labelled NMVs adhered to monocytes (Supplemental Figure 7B) and also NMVs did not activate monocytes (as demonstrated by a lack of L-selectin shedding and no induction of CD11b expression; Supplemental Figure 7C). Thus we conclude that interaction of NMVs released from stimulated neutrophils with endothelial cells, via CD18 on the NMV surface, promotes the subsequent recruitment of monocytes, and that this process does not involve monocyte-NMV interactions. Therefore, we next investigated whether NMVs promote monocyte recruitment by altering endothelial cell inflammatory activation.

### Neutrophil microvesicles induce inflammatory activation of human coronary artery endothelial cells via RelA and increased miR-155 expression

Given the central role of cytokines and adhesion molecules in monocyte recruitment, we investigated the effects of NMVs on endothelial expression of these inflammatory factors. Following 2 h or 4 h incubation with NMVs, HCAEC showed a significant increase in the release of the monocyte chemoattractant CCL2 and the neutrophil chemoattractant CXCL8 compared to HCAEC alone, whereas levels of IL-6 were not significantly increased (Figure 4A). The increase in CCL2 release at 4h is physiologically relevant since it is comparable to that measured when HCAEC were incubated with a well-described inflammatory stimulus, tumor necrosis factor (TNF) (1154 ± 298 pg/mL). In addition, NMVs induced an increase of ICAM-1, vascular cell adhesion molecule-1 (VCAM-1), and CCL2 protein levels and mRNA expression (Figure 4B and C) in HCAEC. These cytokines and adhesion molecules were not detectable in NMVs alone (data not shown) indicating that their production by HCAEC can be induced by NMVs. Importantly, NMVs released by unstimulated neutrophils were unable to induce a significant alteration in gene expression in HCAECs (Supplemental Figure 8A) suggesting that only NMVs released from stimulated neutrophils are able to induce endothelial cell activation.

**Figure 4.**
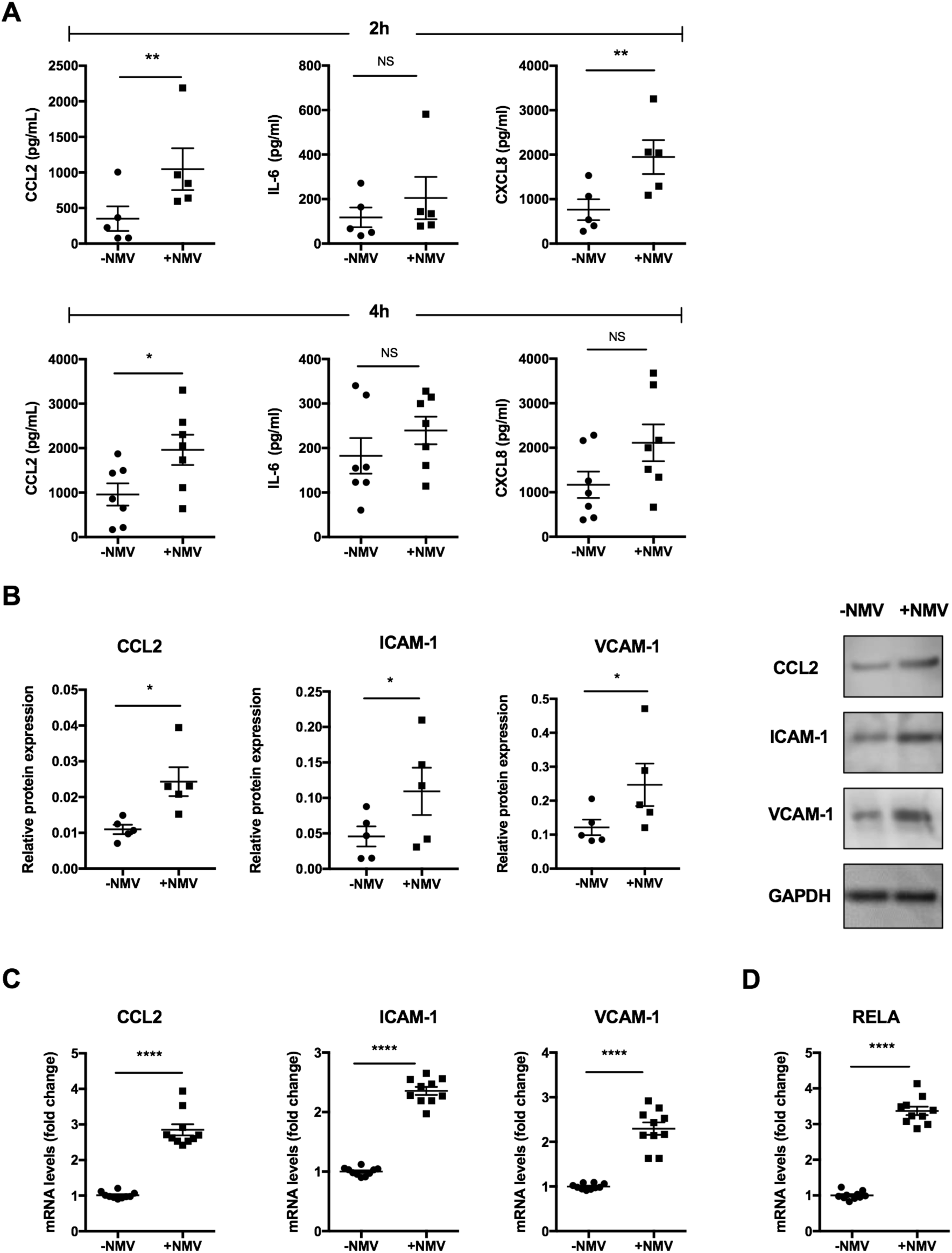
HCAEC activation by neutrophil microvesicles. (A) HCAEC were cultured for 72 h and were then incubated with (+NMV) or without (-NMV) NMVs for 2 h (n = 5) and 4 h (n = 7). Release of CCL2, CXCL8 and IL-6 into the media was analysed using cytometric bead array. (**B - D**) HCAEC were incubated with (+NMV) or without (-NMV) NMVs for 2 h. Alterations in inflammatory protein (**B**; n = 5) and gene (**C**; n = 10) expression were investigated using western blotting and RT-qPCR, respectively. (**D**) *RELA* mRNA levels were measured using RT-qPCR (n = 10). Samples were quantified using densitometry and normalized with β-actin for RT-qPCR and GAPDH for western blot. Results are presented as mean ± SEM and statistical significance evaluated using a paired *t*-test. * *P* < 0.05, ** *P* < 0.01, *** *P* < 0.001.

To interrogate the mechanism by which NMVs induced increased expression of inflammatory molecules, we investigated whether they influence the NF-κB pathway, which is a central regulator of inflammation in arterial endothelial cells. We focused on RELA, a abundant proinflammatory NF-κB subunit in endothelial cells, and found that NMVs released by activated neutrophils induced *RELA* expression in HCAEC after 2 h (Figure 4D) unlike those released by unstimulated cells (Supplemental Figure 8B). We hypothesised that this was mediated via transfer of NMV cargo to endothelial cells and induced inflammatory activation via increased RELA expression. Consistent with this, we found that NMVs were internalized by arterial endothelial cells, both *in vitro* and *in vivo* (Figure 5A - C, Supplemental Movies 1 and 2). This was an active process as internalization was significantly reduced when cells were incubated at 4°C or room temperature compared to 37°C (Figure 5D). Furthermore, TNF, a known inducer of ICAM-1 expression, increased internalization whereas blocking ICAM-1 with anti-ICAM antibody partially inhibited NMV internalization (Figure 5E), both of which support a role for ICAM-1. Internalization *in vivo* was only detected at atheroprone sites and not at atheroprotected sites (Supplemental Movie 3).

**Figure 5.**
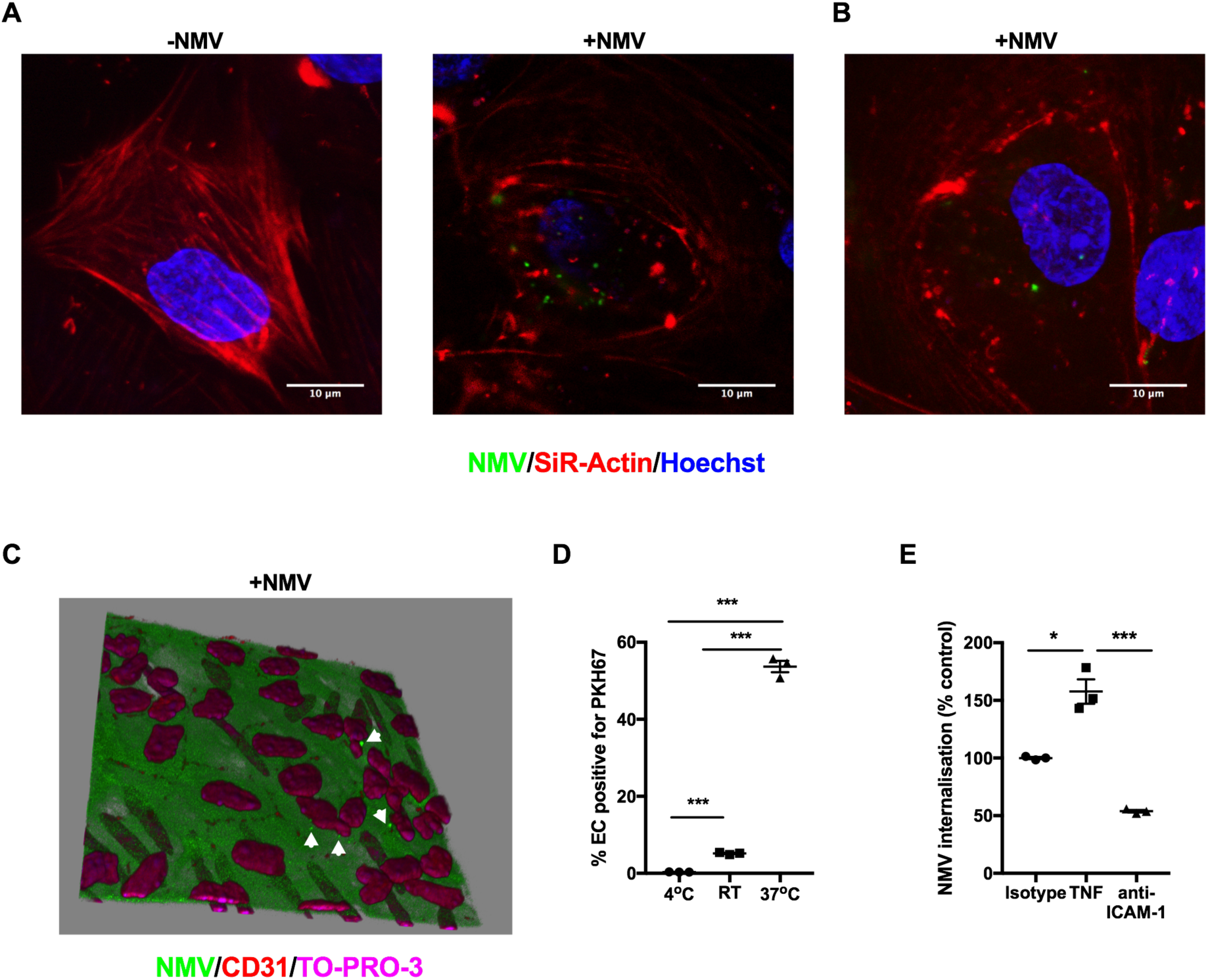
Neutrophil microvesicle internalization by endothelial cells. (**A**) Confocal image and (**B**) Z-stack maximal intensity projection of HCAEC incubated with (+NMV) or without (-NMV) fluorescently-labelled NMVs (green) and stained with SiR-Actin (F-actin; red) and Hoechst (nuclei; blue). (**C**) 3D reconstruction of an *ApoE*^-/-^ mouse aorta stained *en face* with anti-CD31 (endothelial cells; red) and TO-PRO-3 (nuclei; magenta) showing internalization of fluorescently-labelled NMVs (green) 2 h after i.v. injection. Elastin autofluorescence is also shown in green. CD31 expression on the apical surface was used for orientation and the plane of view set just below. Note the misaligned endothelial cell nuclei, characteristic of an area of disturbed flow. Arrows denote NMVs. (**D - E**) HCAEC were incubated with fluorescently-labelled NMVs for 2 h. Fluorescence from residual surface bound NMVs was quenched with trypan blue and data were analysed for changes in mean fluorescence intensity by flow cytometry. (**D**) The experiment was performed at 4°C, room temperature (RT) or 37°C (n = 3). (**E**) HCAEC were incubated at 37°C in the presence of TNF (4 h prior to the addition of NMVs) or anti-ICAM-1 or isotype control (n = 3). Data are expressed as a percentage of the mean of the isotype control samples. Data are presented as mean ± SEM and statistical significance evaluated using one-way ANOVA followed by Tukey’s post test for multiple comparisons. *** *P* < 0.001.

We next investigated whether NMVs contained miRs that could influence gene expression in recipient endothelial cells. RT-qPCR analysis revealed that NMVs contained several miRs, including regulators of inflammation (*miR-9, miR-150, miR-155, miR-186, miR-223*; Figure 6A). We determined whether plasma MV expression of the most abundant miRs, *miR-223* and *miR-155*, were altered by high fat diet in human subjects and found that only *miR-155* expression levels were significantly elevated (Figure 6B). Furthermore, *miR-155* expression levels were found to be increased in plasma MVs and NMVs in mice fed on a Western diet compared to chow (Figure 6C and D). *miR-155* has been shown to increase *NF-κB* expression by targeting its negative regulator, BCL6 ^43–45^. We therefore hypothesised that NMVs activate *NF-κB* by delivering *miR-155*, which reduces *BCL6* expression. Consistent with this, incubation of HCAEC with NMVs led to enhanced endothelial expression of *miR-155* associated with reduced *BCL6* expression (Figure 6E). Futhermore, when HCAEC were incubated with NMVs that were isolated from subjects exposed to high fat diet for 1 week, the increase in *miR-155* expression levels was even greater than that induced by NMVs isolated prior to the diet (Figure 6F). The potential of microvesicles to deliver miR-155 to EC was confirmed by demonstrating that NMVs from wild-type mice can enhance *miR-155* expression in HCAEC whereas NMVs from *miR-155*^-/-^ mice cannot (Figure 6G). Indeed, we found a small but statistically significant reduction in *miR155* copy number in HCAEC when incubated with *miR155*^-/-^ NMVs, although the mechanism for this unexpected finding are unknown and could be due to levels being near the lower detection limit of the assay. The reduced expression of *BCL6* seen with NMV incubation was reversed in endothelial cells transfected with an antagomir that blocks *miR-155* function, indicating that BCL6 is negatively regulated by *miR-155*. Consequently, the increase in expression of *RELA* and its downstream target genes *VCAM-1, ICAM-1* and *CCL2* induced by NMVs was significantly decreased in the presence of *miR-155* antagomir (Figure 6H). Therefore, we deduce that NMVs induce NF-κB activation in endothelial cells via delivery of *miR-155*, which reduces expression of the negative regulator BCL6. In support of this, injection of NMVs into *ApoE*^-/-^ mice induced a significant increase in arterial expression of *miR-155* (Figure 6I) and a subsequent reduction in BCL6 expression (Figure 6 J-K).

**Figure 6.**
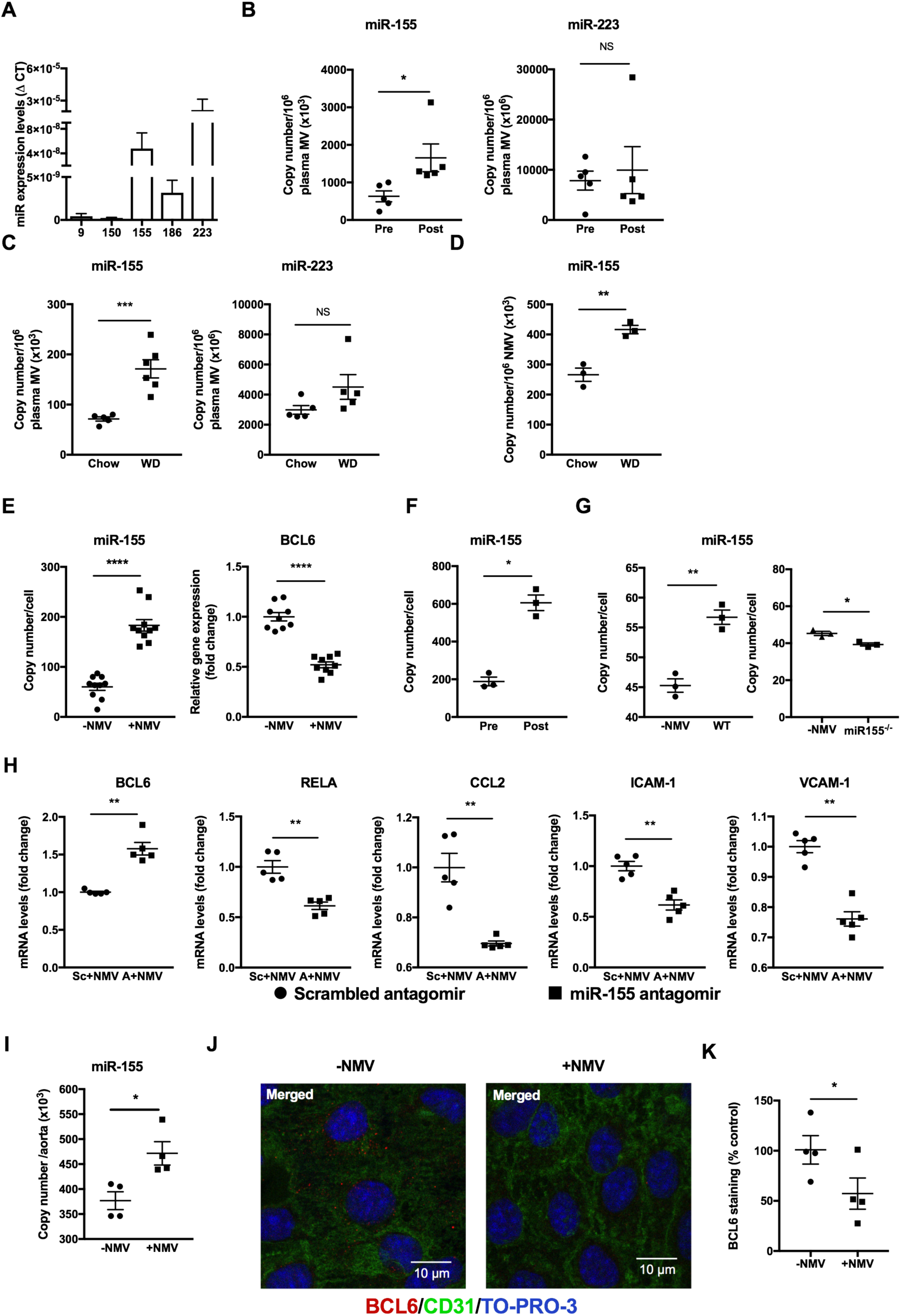
Neutrophil microvesicles contain miRs that are delivered to endothelial cell and alter target gene expression. (**A**) miR content of NMVs prepared from stimulated human neutrophils quantified by RT-qPCR (n = 5). (**B**) *miR-155* and *miR-223* content of MVs in human plasma pre- and post high fat diet (HFD, n = 5), or (**C**) plasma from mice fed chow or Western diet (WD, 6 weeks; n = 5) quantified by RT-qPCR. (**D**) *miR-155* expression levels in NMVs isolated from mice fed chow or Western diet (n = 3) measured by RT-qPCR. (**E**) HCAEC were incubated with (+NMV) or without (-NMV) NMVs for 2 h and *miR-155* and *BCL6* expression levels measured by RT-qPCR (n = 5-10). (**F**) HCAEC were incubated with NMVs prepared from human neutrophils isolated pre- and post HFD and *miR-155* expression levels quantified by RT-qPCR (n = 3) (**G**) HCAEC were incubated with NMVs prepared from *miR-155*^-/-^ vs. wild type mouse neutrophils and *miR-155* expression levels quantified by RT-qPCR (n = 3). (**H**) HCAEC were transfected with 25 ?mol of *miR-155* antagomir (A) or scrambled control (Sc) and incubated with NMVs for 2h. HCAEC expression of *BCL6, RELA* and its downstream targets was investigated by RT-qPCR (n = 5). (**I**) *ApoE*^-/-^ mice were injected i.v. with NMVs via the tail vein and *miR-155* expression levels in mouse the aorta 2h after injection of saline or NMVs (n = 4) was quantified by RT-qPCR. All RT-qPCR samples were normalized with β-actin. (**J - K**) Carotid arteries were isolated from *ApoE*^-/-^ mice and incubated *ex vivo* with (+NMV) or without (-NMV) NMVs for 2 h and stained *en face* with anti-BCL6 antibody (red). Endothelial cells were identified by staining with anti-CD31 (green) and cell nuclei were identified using TO-PRO-3 Iodide (blue). (**J**) Representative *en face* images. Scale bar = 10 µm (**K**) Total fluorescence intensity of BCL6 expression was quantified using ImageJ software (n = 4) and expressed as a percentage of the mean fluorescence in the control samples (-NMV). Data are presented as mean ± SEM and statistical significance evaluated using a paired or unpaired *t*-test as appropriate. * *P* < 0.05, ** *P* < 0.01, *** *P* < 0.001**** *P* < 0.0001.

### Neutrophil microvesicles activate NF-κB and enhance plaque formation in vivo

Having shown regulation of *RelA* expression *in vitro*, we investigated whether circulating NMVs could induce focal inflammation *in vivo*. To investigate this, *ApoE*^-/-^ mice were injected twice weekly for 6 weeks with NMVs to chronically increase circulating NMV levels by approximately 30%, similar to the increase in circulating NMVs observed in human subjects after 7 days on an atherogenic diet (Figure 1D). Levels of RELA in the aorta were assessed by *en face* confocal microscopy and found to be markedly and selectively increased at atheroprone regions in response to NMV injection (Figure 7A and B). Under these conditions, RELA localized partially to the nucleus and a proportion located to the cytoplasm, suggesting that NMVs induce partial activation of NF-κB at atheroprone sites, and also prime endothelial cells for enhanced inflammatory responses by increasing total NF-κB expression.

**Figure 7.**
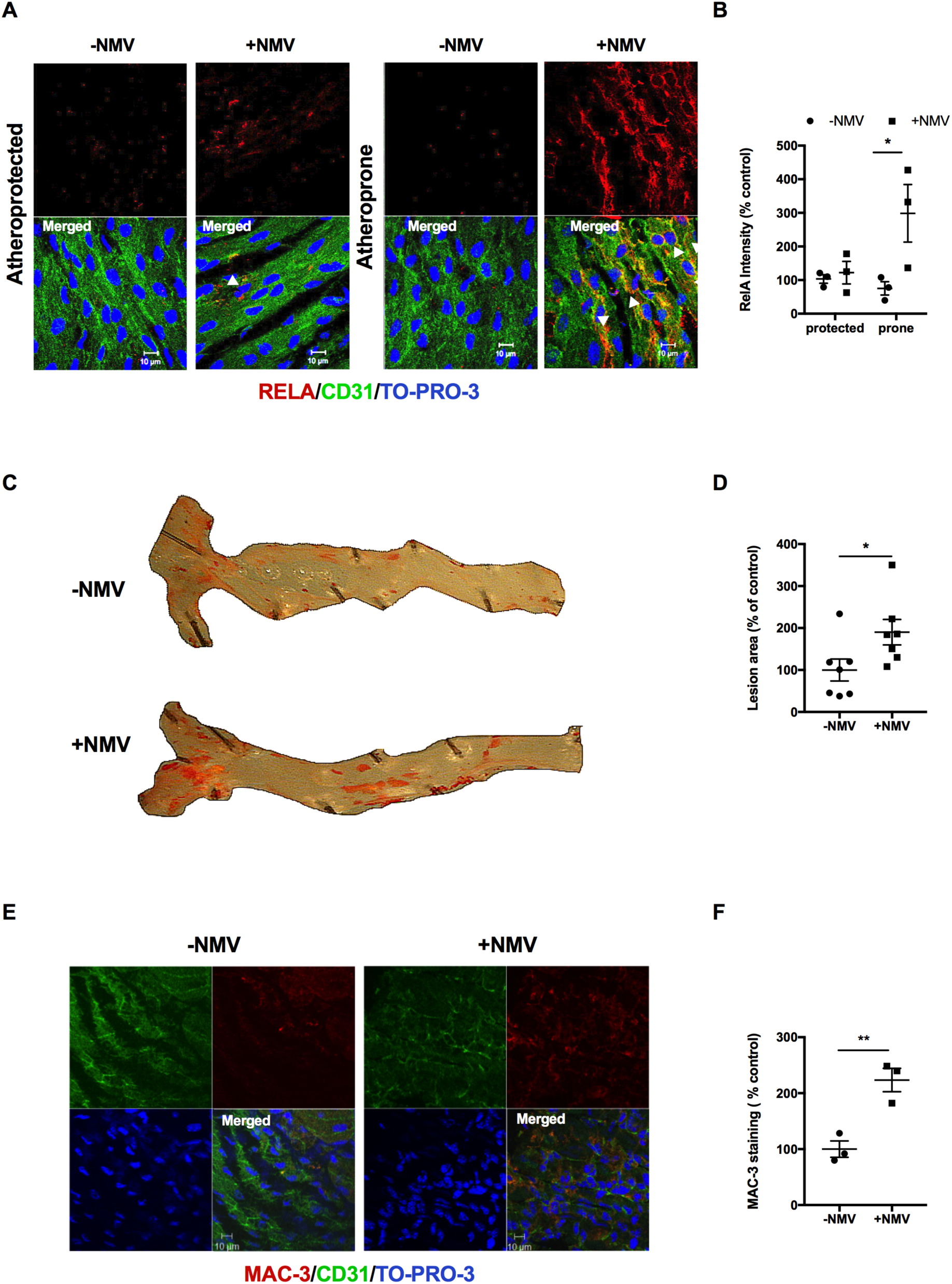
Neutrophil microvesicles induce activation of NF-κB in atheroprone regions and enhance atherosclerosis. *ApoE*^-/-^ mice fed a Western diet were injected with NMVs, or an equivalent volume of saline, twice weekly via the tail vein for 6 weeks. (**A**) Representative *en face* images of RELA (red) expression in descending aorta of mice injected with saline (- NMV) or NMVs (+NMV) in atheroprone regions visualized using confocal fluorescence microscopy. Endothelial cells were identified by staining with anti-CD31 (green) and cell nuclei were identified using TO-PRO-3 Iodide (blue). Examples of nuclear staining of RelA indicated with arrowheads. Scale bar = 10 µm. (**B**) Mean fluorescent intensity was quantified using ImageJ software and data expressed as mean ± SEM (n = 5). (**C - D**) Plaque formation was measured in dissected aortae using *en face* Oil Red O staining and imaged by bright field microscopy. (**C**) Representative images are shown. (**D**) Areas of plaque formation were determined in the entire aorta using NIS-elements analysis software (n = 7). (**E - F**) The aortic arches of mice injected with saline (-NMV) or NMVs (+NMV) were studied by *en face* staining to quantify macrophages (MAC-3, red). (**E**) Endothelial cells were identified by staining with anti-CD31 antibody (green) and cell nuclei were identified using TO-PRO-3 Iodide (blue). Scale bar = 10 µm. (**F**) Mean fluorescence intensity was quantified using Image J software (n = 3). Data are expressed as a percentage of the mean of the control samples (-NMV) and presented as mean ± SEM. Statistical significance was evaluated using two-way ANOVA followed by Bonferroni’s post hoc test (B) or an unpaired *t*- test (D and F). * *P* < 0.05, ** *P* <0.01.

Having determined that NMVs preferentially adhere to atheroprone regions, enhance monocyte transendothelial migration and regulate RELA, we hypothesized that this could lead to enhanced plaque formation. We therefore raised circulating levels of Consistent with this hypothesis, Oil Red O staining revealed significantly more atherosclerotic plaque formation in mice treated with NMVs (over a 6 week period as above) compared to those treated with saline (Figure 7C and D). *En face* staining with MAC3 antibody revealed enhanced recruitment of monocytes / macrophages in response to NMV injection (Figure 7E and F). Thus, we conclude that systemic NMVs can induce focal activation of NF-κB at atheroprone sites and, thereby, amplify vascular inflammation and accelerate lesion formation.

## Discussion

Neutrophils are the most abundant leukocyte in human circulation and are essential for an effective innate immune response. There is also increasing evidence for their role in atherosclerosis. High fat feeding, both in humans and in mouse models, increases the level and activation of neutrophils ^46–48^. Although small numbers of neutrophils have been detected within the core of developing atherosclerotic plaques, this peaks at early stages (4 weeks after high fat diet for *ApoE*^-/-^ and 6 weeks for *LDL*^-/-^ models of experimental atherosclerosis) and neutrophils are rarely observed beyond these time points ^48,49^. Nevertheless, there is evidence to suggest that neutrophils may play a role in plaque development through mechanisms that do not require them to be present within the plaque core, such as neutrophil extracellular trap formation (NETosis) in response to the presence of cholesterol crystals ^50^. In addition to NETosis, activated neutrophils are known to release MVs ^30,35,51^. However, a role for NMVs in atherosclerosis has not previously been investigated. Here we show for the first time that: (i) high fat diet raises levels of circulating NMVs; (ii) NMVs adhere preferentially to atheroprone regions; (iii) once adherent, NMVs are internalized by endothelial cells and deliver *miR-155* and that (iv) NMVs from stimulated neutrophils activate endothelial cells, enhance monocyte recruitment and exacerbate atherosclerotic plaque formation (see Figure 8 for proposed molecular model).

**Figure 8.**
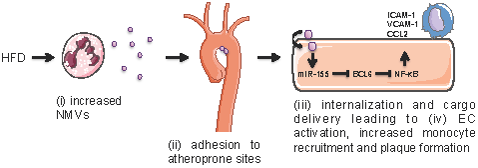
Proposed model of the molecular mechanism of action of neutrophil microvesicles in atherosclerosis.

We observed for the first time that a high fat diet, a known driver of atherosclerosis ^52,53^, potently enhanced circulating levels of NMVs in healthy humans. This was mirrored in hypercholesterolemic mice and led to an accumulation of NMVs in vessel walls. This is in agreement with findings from Leroyer and colleagues who detected the presence of granulocyte-derived MVs in human atherosclerotic plaques ^18^. As well as NMVs, platelet and monocyte MVs also accumulated in the vessel wall and it is possible that the accumulation of one MV subpopulation can influence the production and recruitment of other MVs. As well as NMVs, platelet and monocyte MVs undoubtedly contribute to the local microenvironment within the plaque and detailed mechanistic studies into the role of these specific MVs in vascular inflammation and atherosclerotic plaque development are an exciting area for future research.

Atherosclerosis is a focal disease that occurs in regions of disturbed flow. NMV-endothelial cell interactions were increased when arterial cells were conditioned under disturbed flow and at atheroprone sites in mice fed on a high fat diet. This correlated with the expression of ICAM-1, and adhesion was indeed found to be ICAM-1 dependent. Interestingly, interaction of NMVs from stimulated neutrophils with endothelial cells further increased ICAM-1 expression suggesting that this interaction may lead to an augmentation of subsequent NMV-endothelial cell interactions (i.e. positive feedback). This could provide a mechanism by which NMVs accumulate in the vessel wall of mice on a high fat diet over time. Once adherent, NMVs were internalized by endothelial cells. ICAM-1 was also required for the internalization of NMVs and, interestingly, Muro *et al.* have described a novel pathway for endocytosis involving clustering of ICAM-1 ^54,55^. It is plausible that NMVs utilize this pathway, providing a mechanism by which increased levels of internalization occur in areas where there is increased expression of this adhesion molecule, such as atheroprone regions. Further studies to investigate the precise mechanisms by which NMVs are internalized are of great importance as this could potentially be exploited in the future for targeted delivery of therapeutic agents.

Although we found that unstimulated neutrophils released NMVs, we also found that, unlike NMVs from stimulated neutrophils, these NMVs were unable to induce endothelial cell activation. It is likely that these functional differences are due to divergent cargo as previously described for NMVs released from adherent vs. non-adherent neutrophils ^56^. There has been much interest in the ability of extracellular vesicles to transfer genetic material from the parent cell to a target cell ^14,15^. Here we demonstrate that NMVs contain numerous miRs implicated in atherosclerosis and vascular inflammation, the most abundant of which were *miR-155* and *miR-223*. High fat feeding of both healthy human volunteers and *ApoE*^-/-^ mice resulted in a significant increase in plasma MV and NMV levels of *miR-155* but not *miR-223*. In addition, incubation of HCAECs with NMVs increased cellular expression of *miR-155*, which was further increased when NMVs were prepared from subjects who had undertaken the high fat diet study. We concluded that miR-155 was delivered to, but not induced in, HCAEC as NMVs isolated from *miR-155*^-/-^ mice did not enhance *miR-155* expression in HCAEC. We subsequently carried out functional studies using a miR-155-specific antagomir and demonstrated that miR-155 enhanced inflammatory gene expression by promoting the expression of the *RELA NF-κB* sub-unit. The mechanism involves BCL6, a negative regulator of NF-κB, whose expression in EC was suppressed by miR-155 delivery. Thus, we suggest that NMVs enhance NF-κB expression in endothelial cells by delivering miR-155, which inhibits the negative regulator BCL6. Consistent with this, we observed that NMVs enhance NF-κB expression at atheroprone regions in hypercholesterolemic mice. Moreover, *miR-155* expression is increased in both human and mouse atherosclerotic plaques, and *ApoE*^-/-^ mice deficient in *miR-155* have reduced atherogenesis and monocyte recruitment ^57^. Thus, we hypothesize that NMVs play a role in focal increases in miR-155 levels, leading to enhanced vascular inflammation and accelerated atherogenesis. It should be noted that NMVs contain a complex mixture of proteins, RNAs, and miRs that may also contribute to atherosclerosis development and future studies should address the potential role of these factors. Nevertheless, despite this complex cargo, we have shown that miR-155 is an important component and may be a potential therapeutic target in atherosclerosis.

Together, our studies presented here provide novel, fundamental insights into the mechanism of action of NMVs in enhancing vascular inflammation and monocyte recruitment to developing plaques, potentially solving a long-standing enigma regarding the role of neutrophils in atherosclerosis. The ability of NMVs to preferentially adhere to atheroprone sites and activate endothelial cells could be a major mechanism by which neutrophils contribute to plaque formation but are rarely detected in human plaques.

## Acknowledgments

We thank Dr F. Stassen for the use of the qNano Gold and K. Knoops for advice using the Amira software.

## Contribution of Authors

IG, BW, CS, MA and CR contributed to the design of experiments, acquisition and analysis of data and preparation of the manuscript. JJ, AB, MM, LAL, LW, ML, SP, RW CH and MZ contributed to the acquisition and analysis of data. LW, BB, RB, SF, EKT and AS contributed to the acquisition of data. PH contributed to the initial design of the study and reviewed the manuscript. PE contributed to the conception and design of experiments, the analysis of data and the preparation of the manuscript. VR conceived the study, contributed to the acquisition of data, analysed the data and wrote the manuscript.

## Non-standard Abbreviations and Acronyms

*ApoE*^-/-^: apolipoprotein E deficient
HCAEC: human coronary artery endothelial cells (HCAEC)
ICAM-1: intercellular adhesion molecule-1
miRs: micro RNA
MVs: microvesicles
NETosis: neutrophil exracellular trap formation
NF-κB: nuclear factor
NMVs: neutrophil-derived microvesicles
OSS: oscillatory shear stress
qPCR: quantitative polymerase chain reaction
RT-PCR: reverse transcription polymerase chain reaction
REE: resting energy expenditure
TNF: tumor necrosis factor
TRPS: tunable Resistive Pulse Sensing
VCAM-1: vascular cell adhesion molecule-1

